# Micro-CT: A reversible contrast-based protocol for non-destructive imaging of cardiac morphology in an avian model

**DOI:** 10.64898/2026.07.15.737570

**Authors:** Joshua Durrans, Nicola Aberdein, Prachi Stafford, Liam Ridge, Mari Herigstad

## Abstract

Microcomputed tomography (micro-CT) is a useful tool that can be utilised for 3D structural characterisation and volumetric quantification of small biological specimens. Its potential application is particularly valuable within the field of cardiac development, where phenotypic profiling at the whole organ, cell, and molecular level is often most informative within the same sample. Consequently, this study sought to develop a multimodal imaging protocol to enable 3D phenotypic characterisation of embryonic avian hearts (iodine-based contrast X-ray imaging) prior to immunohistochemistry-based cell and molecular analysis. Micro-CT parameters were tested to establish an optimal protocol for 3D analysis of embryonic cardiac specimens across multiple developmental timepoints. Optimised parameters provided reliable and reproducible 3D analysis of cardiac macrostructures. Sodium thiosulphate treatment of X-ray imaged hearts effectively reversed the iodine-based contrast stain whilst maintaining antigen availability of nuclear, membranous, and cytoplasmic targets in traditional downstream imaging studies. Together, this study demonstrates a robust and highly efficient multimodal imaging strategy to comprehensively characterise cardiac morphology in avian embryos and may serve as a versatile foundation for a broad range of bioimaging applications within the wider scientific community.

## INTRODUCTION

### Characterisation of congenital heart defects in biomedical research

Comprehensive cardiac developmental studies into congenital heart defects (CHD), a significant cause of mortality and morbidity throughout the world (Hoffman & Kaplan, 2002), often require both whole-organ morphology and molecular-level information. Imaging techniques such as hybridisation chain reaction and light sheet microscopy are valuable spatial molecular profiling techniques, but do not fully capture simultaneous structural changes (Degenhardt et al., 2010; Matias et al. 2025; Metscher, 2009) required to characterise and interpret the mechanistic alterations underpinning CHDs (Liu et al., 2014; Schulte et al., 2024; Weber & Huisken, 2011). Moreover, they cannot reliably quantify whole-organ geometry or luminal volume, measurements that provide insight into potential physiological changes that disrupt cardiac output and overall functionality (Kyhl et al., 2013). Microcomputed tomography (micro-CT) can bridge this critical gap by enabling analysis of whole-organ cardiac morphology, identifying subtle perturbations in chamber geometry, myocardial architecture and septal structure that underpin many CHDs. CHD aetiology involves both environmental and genetic factors which often results in incomplete penetrance (Akhirome et al., 2017); therefore, a comprehensive in-depth phenotypic assessment is vital. Developing an approach that preserves a specimen for additional cell and molecular analysis would provide substantial value by enabling multi-layered analysis of the same biological specimen.

Traditional FFPE-based immunohistochemistry remains the gold-standard approach to visualise protein localisation within tissue specimens (Bauman et al., 2016); however, this technique is inherently destructive and renders the specimen unsuitable for additional experimentation (Khimchenko et al, 2016; Zhang et al., 2022). In addition, common operator limitations include a loss of continuous tissue sections and introduction of artifacts, such as tissue folding, which can affect imaging continuity and prevent accurate morphological interpretation (McInnes, 2005). Furthermore, ventricular morphology is highly regionalised with myocardial thickness, trabeculae density and septal structure varying across regions within the heart; this limits mechanistic interpretation, particularly during CHD pathophysiology, due to incomplete penetrance within individual embryos and knock-out models varying in the presence or severity of the structural defects being investigated (Akhirome et al., 2017). This reinforces the need for a multimodal workflow that captures whole-organ morphology before targeted molecular analysis.

### Integration of micro-CT imaging into a multimodal workflow

Micro-CT is a high-resolution imaging technique that can provide high-spatial resolution down to an effective voxel size of 1μm and allow visualisation of both macro- and fine cardiac structures such as the heart chambers, outflow tract, valve leaflets and myocardial trabeculae in a three-dimensional (3D) space (Keklikoglou et al., 2021). This provides valuable information which would assist in the characterisation of CHDs and provide an additional quantitative analysis tool to non-destructively visualise the internal architecture of intact cardiac tissue (Campbell & Sophocleous, 2014; Rühli et al., 2007). Although other non-destructive approaches exist, such as magnetic resonance imaging (MRI), optical coherence tomography and ultrasound, these are more useful for larger samples as they offer lower spatial resolution with poorer contrast between cardiac morphology and intralumenal space, or are unsuitable for the reduced specimen size of embryonic tissue in smaller organisms such as mice and chicken embryos (Ruffins et al., 2007). Although ultra-high-field MRI imaging (>9T) can image small *ex-vivo* specimens, micro-CT provides substantially higher spatial resolution (<1μ voxel size, compared to <200μ in UHF MRI) for embryonic chicken hearts (2-5mm in diameter, compared to 8-9cm of an adult human heart), enabling detailed assessment of structural cardiac morphology (Ivanov et al., 2022).

Confocal microscopy is a powerful destructive imaging tool that provides high-resolution sub-cellular localisation of biological molecules (Jonkman & Brown, 2015) that cannot be achieved with non-destructive modalities such as micro-CT (Metscher, 2009). Its ability to localise the expression of signalling molecules enables mechanistic interpretation of cellular pathologies (Jonkman & Brown, 2015). However, confocal imaging requires optical clearing of larger samples (>200μm thickness), or sectioning which disrupts or destroys native 3D geometry and restricts analysis to small regionally limited volumes (Clendenon et al., 2015; Makki et al., 2025). This compounds the challenge of analysing subtle, regional-specific differences in cardiac morphology whilst also excluding these biological specimens from other experimental applications, which is particularly important when working with variably penetrant cardiac phenotypes where every specimen is valuable (Clendenon et al., 2015; Jonkman & Brown, 2015). Confocal imaging also requires substantial labour-intensive sample preparation steps which include fixation, dehydration, clearing, rehydration and staining steps that takes several days to complete (Jonkman & Brown, 2015; Makki et al., 2025). This limits the number of specimens that can be processed and analysed at any one time.

The labour-intensive preparation required for confocal imaging (Jonkman & Brown, 2015; Makki et al., 2025) contrasts significantly with the rapid and minimal sample preparation required for micro-CT imaging (Metscher, 2009). This further reinforces the value of a sequential multimodal workflow which includes rapid non-destructive imaging of whole-organ morphology followed by targeted confocal immunohistochemistry to be performed only where it is the most informative, increasing resource-efficiency and maximises the information gained from each valuable specimen.

### Developing a reversible contrast strategy for a dual micro-CT and immunohistochemistry approach

Due to being an endogenous source of contrast in biological specimens, micro-CT is widely used to visualise high density mineralised tissue such as bone (Lindtner et al., 2025). A disadvantage of imaging soft tissue such as the heart, is the lack of contrast between macro-structures such as the myocardium and luminal space which may be filled with fixative resulting in no clear differentiation between the structures (Metscher, 2009; Procházková et al., 2011). To circumvent this, strategies have been developed to artificially enhance specimen contrast prior to imaging (de Bournonville et al., 2019). There are many types of contrast-enhancement dyes for micro-CT, including phosphotungstic acid, phosphomolybdic acid and osmium tetroxide (Pauwels et al., 2013). These contrast agents yield high-resolution imaging data but require strong destaining techniques for further downstream applications as they bind strongly with collagen, proteins, and lipids (Metscher, 2009). The additional destaining steps required can disrupt tissue integrity and antigen availability, and manifest in subsequent imaging artefacts (Gignac et al., 2016; Lanzetti & Ekdale, 2021; Metscher, 2009). This is illustrated by use of phosphotungstic acid, which has previously been shown to show good contrast in soft-tissue but, its acidic nature compromises tissue integrity by inducing soft-tissue shrinkage (de Bournonville et al., 2019). For a multimodal approach to work, a contrast dye that has minimal effect on antigen availability must be used.

Lugol’s stain has been demonstrated to enhance contrast whilst being an easily reversible contrast-dye using sodium thiosulphate (Hopkins et al., 2015). Whilst Lugol’s is superior to other enhancement dyes in this regard, it is important to note that sodium thiosulphate destaining does not restore a biological specimen to its original state and that iodide remains within the tissue (Gignac et al., 2016). Whilst Lugol’s has been shown to cause substantial soft-tissue shrinkage (Heimel et al., 2019; Vickerton et al., 2013), this can be reduced by using a buffered Lugol’s solution. Dawood et al. (2021) identified that shrinkage is caused by pH changes and that mouse livers stained in a Sorensen’s buffered Lugol’s solution resulted in a shrinkage of 5.9%, compared to 31.9% when not using a buffered stain. With this in mind, buffered Lugol’s iodine has emerged as a promising contrast agent for establishing this multimodal workflow, offering rapid, high-contrast micro-CT staining without compromising subsequent confocal immunohistochemistry.

### Aims and Objectives

This study aimed to determine whether existing contrast-enhanced micro-CT protocols using Lugol’s iodine (Dawood et al., 2021; Metscher, 2009) could be adopted into the traditional immunohistochemistry-based workflow. Micro-CT was chosen as the first modality due to it being a non-destructive technique that requires minimal sample preparation and provides fast image acquisition to provide visualisation of whole-organ structures. The second modality was confocal imaging after sodium thiosulphate destaining and destructive FFPE sectioning to complement structural phenotyping with cellular-level molecular analysis.

The first aim was to optimise the iodine-based micro-CT staining protocol followed by parameter selection to optimise high-resolution images across multiple developmental timepoints during cardiogenesis to characterise cardiac phenotypes. The second aim was to establish whether further downstream imaging is possible by probing for cytoplasmic (β-actin), nuclear (phosphohistone-H3), and membrane (WGA) targets after sample destaining to confirm antigen availability and validate the dual multimodal workflow.

## MATERIALS AND METHODS

### Ethical Approval

Ethical approval was granted by the university research ethics committee at Sheffield Hallam University (Ethics Review number ER64386858), and all experiments were undertaken in accordance with the rules and regulations outlined by the UK Home Office and Animals (Scientific Procedures) Act 1986.

### Egg Incubation

Fertilised Shaver Brown chicken eggs (*Gallus gallus domesticus*) were used for all experiments (MedEggs Ltd., UK). The eggs were incubated in a humidified incubator at 37.5°C and turned 180° each day at three timepoints (at 9:00am, 12:30pm and 5:00pm) to prevent the embryo from sticking to the shell membrane. On the required day of dissection (embryonic day (D)7, D10 or D12), the eggs were removed from the incubator, and each egg was opened with forceps. The embryo was moved to a petri-dish containing ice-cold PBS (Gibco, 14040133) and assessed for viability (presence of heartbeat). All nonviable embryos were excluded. After decapitation, hearts were excised by severing the outflow tract/great vessels and fixed in 4% paraformaldehyde for 30 minutes before being washed in PBS (3 x 30 minutes). After fixation, the hearts were blotted dry on tissue paper and weighed before being placed in fresh PBS and stored at 4°C until processed for micro-CT imaging.

### Lugol’s Staining

Samples were stained in a 2% Lugol’s solution prepared in 1X Sorensen’s phosphate buffer (133 mM total phosphate, pH 7.2). 2X Sorensen’s was prepared at 266 mM total phosphate (pH 7.2). Lugol’s stock solution was 15% (w/v), containing 5% iodine and 10% potassium iodide as described in Dawood et al. (2021).

The embryonic heart was placed in a glass sample vial containing 5mL of the staining solution. The glass vial was placed on an orbital shaker at 4°C for 24 hours prior to micro-CT imaging. On the day of imaging, the stained sample was briefly rinsed in PBS at room temperature to remove excess iodine from the heart chambers.

### Sample Mounting and Scan Acquisition

Each biological specimen was positioned inside a PCR tube (Starlab, I1402-8100) with a minimal volume (20-40µL) of PBS to maintain sample hydration and prevent tissue shrinkage during µCT scanning. The tube was inverted and placed on the instrument stage of a Bruker SkyScan 1272 (Bruker, United States) as illustrated in Figure 1.

**Figure 1.**
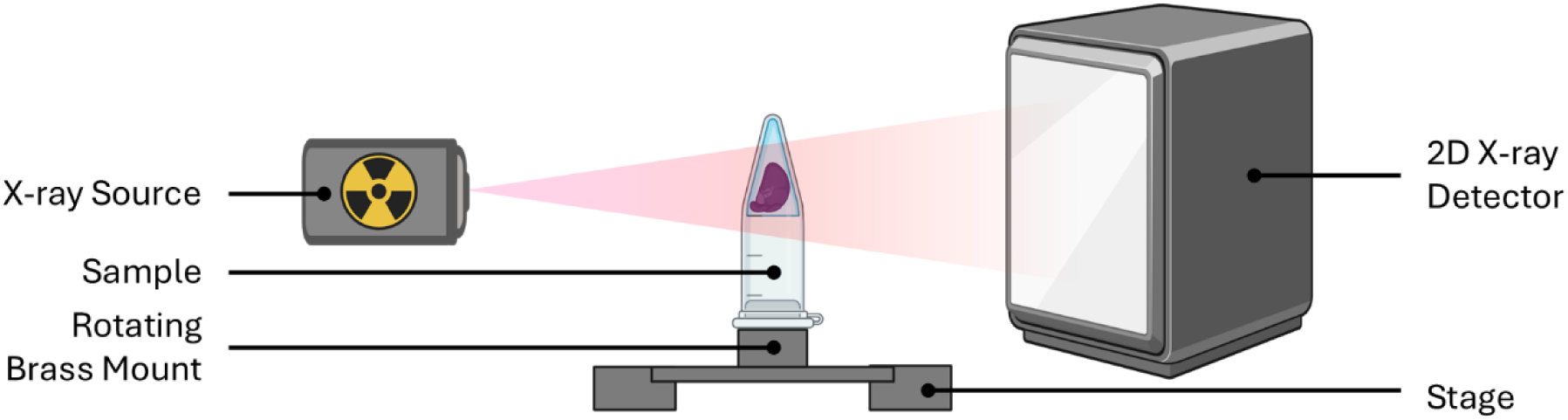
Schematic of sample mounting. Illustration of the Bruker Skyscan 1272 mounting setup.

Micro-CT scans of embryonic chicken hearts were acquired using the Bruker SkyScan 1272 bench-top scanner. The software was controlled using the SkyScan 1272 Control Program (Bruker, United States, Version 1.5.0.0). During the optimisation process, the following parameters were adjusted to identify which settings yielded the most detailed visualisation of tissue: voxel size, filter, camera binning, rotation step and frame averaging as shown in Table 1.

**Table 1.** Parameters used during the optimisation of scans.

|  |  |  |  |  |
| --- | --- | --- | --- | --- |
| **Isotropic Voxel Size** | 3 µm | 4 µm | 5 µm | 6 µm |
| **Filter Selection** | No Filter | Aluminium 0.25mm | Aluminium 0.5mm | Aluminium 1mm |
| **Camera Binning** | 1024x1024 (1K) | 2048x2048 (2K) | 4096x4096 (4K) |  |
| **Rotation Step** | 0.1° | 0.3° | 0.5° | 0.7° |
| **Frame Averaging** | 1 | 3 | 5 |  |

### Reconstruction

Projection data was reconstructed using NRecon (Bruker, United States, Version 2.2.0.6) prior to analysis. The following parameters were used during the reconstruction process: beam hardening correction, ring artefact reduction, and automatic misalignment correction. A gaussian smoothing of 1 was applied during the reconstruction process, additional smoothing operations were applied during post-processing operations using Dragonfly as detailed further below (Comet Technologies, Canada, Version 2024.1 Build 1627).

DataViewer software (Bruker, United States, Version 1.5.6.2) was used to re-orientate scanned samples into a frontal (coronal) alignment for consistent comparisons across specimens.

### Segmentation and 3D Analysis

Reconstructed images were imported into Dragonfly. Image spacing was set to the voxel size reported by the SkyScan 1272. Analyses were saved as Dragonfly session files (.ORSSession) for each individual specimen.

Dragonfly’s Image Processing panel was used to create an operations list that was applied identically to all volumes prior to segmentation and analysis. The workflow comprised of a 3D median smoothing filter (kernel; 3x3x3) to supress background noise whilst preserving cardiac morphology, followed by a 2D closing operation (kernel; 3).

Regions of interest (ROI) were drawn to segment areas for volumetric analysis. Post segmentation, the ROIs were refined to remove non-connected objects from the lumen and wall ROIs. For the wall ROIs, the largest 3D component was retained and objects under 6-connected objects (face-connected) were removed. For the wall ROI, the largest 3D component was retained and objects under 26-connected objects (face-edge-vertex) were removed. The regions of interest were segmented using anatomical landmarks described by Butcher et al. (2006) and Young et al. (2000).

***Left Ventricular Wall* –** Left ventricular wall volume (Fig 2. Red) was defined as the compact and trabecular myocardium comprising the left free wall and the compact interventricular septum and septal trabeculae that extend into the endocardial space of the left ventricle. Analysis extended from the ventricular apex to the ventricular margin of the endocardial cushions of the mitral (bicuspid) valve.

**Figure 2.**
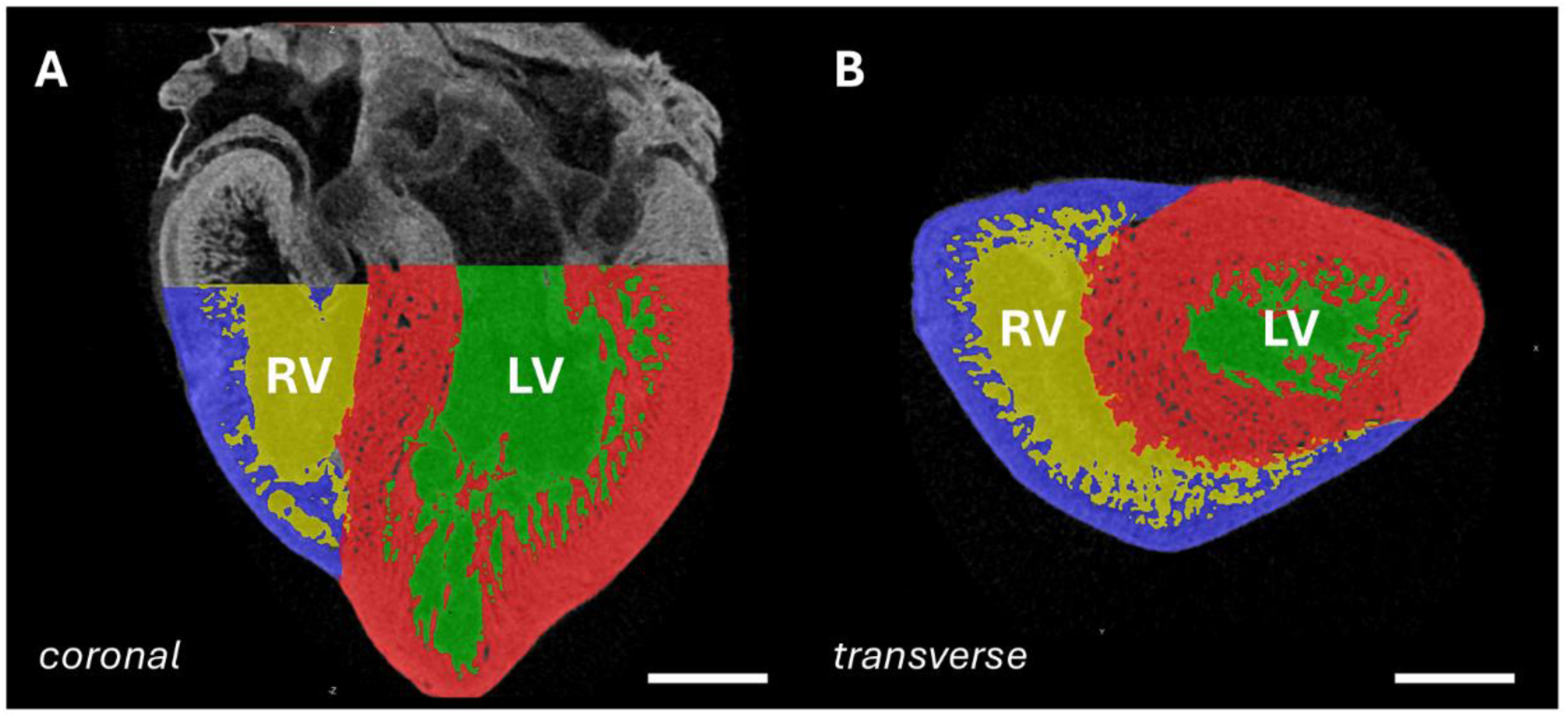
View of image segmentation. **A-B:** View of (**A**) coronal and (**B**) transverse cross-sections of the D10 embryonic chicken heart were segmented into the left ventricular free-wall and interventricular septum (red), right ventricular free-wall (blue), left ventricular lumen (purple), and right ventricular lumen (green) using Dragonfly (CometTech, Canada). Abbreviations: LV-left ventricle, RV-right ventricle. Scale bar is 500μm.

***Left Ventricular Lumen*** – Left ventricular lumen volume (Fig 2. Green) was defined as the endocardial space enclosed between the compact and trabecular myocardium of the left free-wall and the compact septal myocardium and septal trabecular myocardium that extend into the endocardial space of the left ventricle. Analyses extended from the ventricular apex to the ventricular margin of the endocardial cushions of the mitral valve.

***Right Ventricular Wall*** – Right ventricular wall volume (Fig 2. Blue) was defined as the compact and trabecular myocardium comprising the right free wall and the septal trabeculae that extend into the endocardial space of the right ventricle. Analyses extended from the ventricular apex to the ventricular margin of the endocardial cushions of the tricuspid valve.

***Right Ventricular Lumen*** – Right ventricular lumen volume (Fig 2. Yellow) was defined as the endocardial space enclosed between the compact and trabecular myocardium of the right free-wall and the septal trabecular myocardium that extend into the endocardial space of the right ventricle. Analyses extended from the ventricular apex to the ventricular margin of the endocardial cushions of the tricuspid valve.

### Sodium Thiosulphate Destaining and Paraffin Embedding

Immediately post-scanning, specimens were incubated in 2.5% (w/v) sodium thiosulphate (Thermo Scientific, 450620010) for 24 hours at 4°C on an oscillating see-saw rocker and incubated in PBS for a further 72 hours prior to the paraffin embedding.

Specimens were dehydrated in an ethanol series (50, 70, 85, 95, 100%, in PBS) and cleared in Histoclear (SLS, NAT1330) prior to incubation in a 1:1 (v/v) paraffin/Histoclear mix at 65°C with regular changes. Hearts were then cleared in 100% (v/v) paraffin with regular changes as described by Ridge et al. (2021). Specimens were embedded in a base mould and sectioned in coronal orientation at 10μ using a Leica RM2235 microtome (Leica, Newcastle, UK).

### Histopathology and Immunohistochemistry

For H&E staining, slides were de-waxed in Histoclear (National Diagnostics, HS-200) (2 x 10 minutes) and rehydrated in an ethanol series, (100, 70, 50, 30%, in dH_2_O, 5 minutes each) and washed in water. Slides were then incubated in Harris’ haematoxylin (Sigma-Aldrich, HHS32) for 3.5 minutes, washed, and incubated in eosin (Sigma-Aldrich, HT110132) for 90 seconds prior to rinsing and mounting coverslips with DPX (Sigma-Aldrich, 1005790500).

For immunohistochemistry staining, slides were de-waxed in Histoclear (2 x 10 minutes) and rehydrated in an ethanol series, (100, 70, 50, 30%, in PBS, 1 x 5 mins) and washed in PBS (2 x 5 minutes). Antigen retrieval was performed in Sodium Citrate buffer (10 mM trisodium citrate, 0.05% (v/v) Triton X-100, pH 6.0 in 1000ml dH_2_O). Slides were heated for 10 minutes in a laboratory microwave at medium power (380W) and left to cool for 30 minutes. Slides were rinsed in TBS (20mM Tris Base, 150mM NaCl, in 1000ml dH_2_O) and blocked (5% v/v normal Goat serum, in TBS) for 1 hour. Slides were incubated in rabbit anti-phosphohistone-H3 (Invitrogen, PA5-17869, 1:200) or rabbit anti-β-actin (Proteintech, 81115-1-RR, 1:500) primary antibody (diluted in 1% w/v BSA in PBS) and incubated overnight at 4°C. The following day, slides were washed in PBS (2 x 5 minutes) and further incubated in a goat anti-Rabbit Alexa Fluor Plus 488 (Invitrogen, A32731, 1:500) secondary antibody diluted in dilution buffer for 1 hour. Slides were then rinsed in PBS (2 x 5 minutes), incubated in wheat germ agglutinin-conjugated Alexa Fluor 647 (Invitrogen, W32466, 5μg/mL) for 2 minutes, washed in PBS and further incubated in DAPI (Invitrogen, D21490, 1µg/mL) for 2 minutes.

Slides were briefly washed and counter-stained with Sudan Black B (0.1% w/v, in 70% ethanol) (Fisher Scientific, 15494669) for 5 minutes to remove tissue autofluorescence, washed and mounted with Invitrogen Prolong Diamond Antifade (Invitrogen, P36970) prior to microscopic imaging using a Zeiss LSM800 confocal microscope.

### Descriptive Statistics

Mean and standard deviation are shown in text and figure legends as (Mean±SD). Descriptive analysis was performed in Graphpad PRISM 10 (v10.6.1).

Data was normalised by adapting previously published protocols. The volume of the region of interest was divided by heart weight to create a normalised value relative to each individual specimen (Markel et al., 2020; Lloyd et al., 2011).

## RESULTS AND DISCUSSION

This study set out to optimise and develop a multimodal imaging approach for micro-CT phenotyping and morphometric volume analysis to characterise cardiac morphology using reversible iodine-based contrast staining protocol to allow integration into the traditional pathology study of cellular morphology in a by immunohistochemistry, histology, and wider fluorescent microscopy staining techniques and facilitate efficient, comprehensive characterisation of cardiac tissue architecture.

### Filter selection for optimal attenuation

Low energy X-rays are preferentially absorbed by materials whereas high-energy X-rays pass through materials more readily. This creates a heterogenous distribution of X-rays as they pass through a sample leading to beam hardening. To determine which X-ray beam filter was appropriate to reduce beam hardening artefacts and improve image quality, four filter options were tested, as shown in Figure 3. Filter selection is dependent on the composition of the material being scanned, the size and the shape (Bouxsein et al., 2010). Filters placed in front of the X-ray beam narrow the X-ray energy spectrum; this increases the average X-ray energy as the low-energy X-rays are absorbed, subsequently reducing beam hardening and producing a more uniform result. The optimum minimum and maximum x-ray attenuation through a sample should be approximately 30% minimum and 95% maximum to reduce beam hardening artefacts (Bouxsein et al., 2010), To obtain a high-quality scan, it is important to use as much of the full dynamic range of the detector as possible (du Plessis et al., 2017).

**Figure 3.**
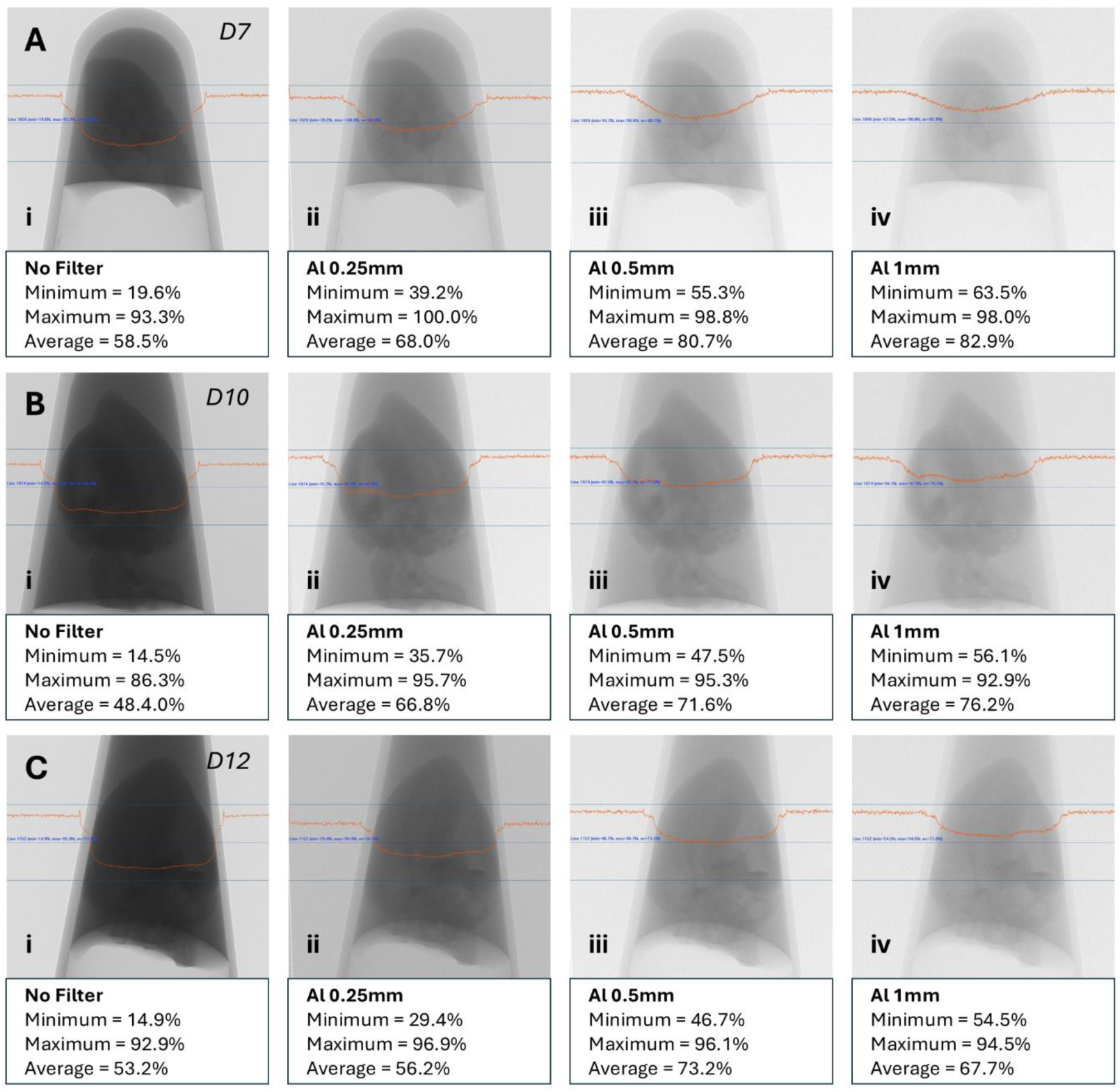
Attenuation of X-rays across biological specimens at different biological timepoints. The optimal attenuation was defined by scanning parameters close to a minimum attenuation of 30%, maximum attenuation of 95% and average attenuation of 70%. **A:** For day 7 (D7), the optimal attenuation was (**i**) no filter to obtain a minimum of 19.6%, maximum of 93.3% and average of 58.5%. For (**B**) day 10 (D10) and (**C**) day 12 (D12), the optimal filter was (**ii**) the aluminium 0.25mm filter to obtain a minimum attenuation of 35.7%, maximum of 95.7% and average of 66.8% for day 10 and minimum of 29.4%, maximum of 96.9% and average attenuation of 56.2% for day 12.

It was determined that for embryonic day 7 samples (Fig 3. A), the optimal filter for an attenuation between recommended 30% and 95% was no filter as the unfiltered images showed higher signal-to-noise, clear tissue boundaries, and no evidence of detector saturation after reconstruction. It was further determined that for embryonic day 10 and 12 (Fig 3. B-C), the optimal filter was an aluminium 0.25mm filter to reduce attenuation into the 30% to 95% range.

### Average number of frames and rotation steps

The average number of radiographs acquired at each rotational point during the micro-CT scanning process should be considered. Increasing the rotational step reduced the total scanning time, whereas increasing the number of radiographs taken increased scanning time. Therefore, it was important to balance the number of repeated radiographs with the rotational step between radiographs to obtain high-resolution images. For iodine-enhanced specimens, minimising scanning time is beneficial, as iodine can leach from the sample into the surrounding PBS which can reduce sample contrast and affect X-ray attenuation and impact reconstruction quality. Contrast enhancement using iodine is also beneficial as it is possible to produce high-quality micro-CT images after a short staining incubation step followed by a short scan duration (Souza e Silva et al., 2017) as other stains such as phosphotungstic acid require a prolonged 7-day incubation depending on sample size (Metscher, 2009).

Increasing the average number of frames reduces signal-to-noise during reconstruction (Rutty et al., 2013). However, an excessive number of average frames does not increase overall scan quality and instead increases the duration of the total scan time (Irie et al., 2022). This was a consideration when studying biological specimens that have been artificially enhanced by a contrast-dye. A rotational step of 0.5° and an average of 3 repeated radiographs at each rotational point generated a high-resolution image suitable for further downstream quantitative analysis. There was no observed benefit to increasing the average number of frames to 5 or 7. It was also determined whether the sample should be scanned for a full 360° rotation or whether a 180° rotation was satisfactory for a scan. A 360° scan doubled the scanning time and there was no difference in scan quality.

### Isotropic voxel size and camera binning

During acquisition, it is crucial to consider the isotropic voxel size as this directly affects the spatial resolution used during downstream quantitative analysis. Smaller isotropic voxels (higher resolution) provide higher precision and greater delineation of fine structures such as myocardial trabeculae and the microvasculature as shown in Figure 4. Camera binning combines adjacent pixels to improve the background signal-to-noise ratio and reduce total scanning time.

**Figure 4.**
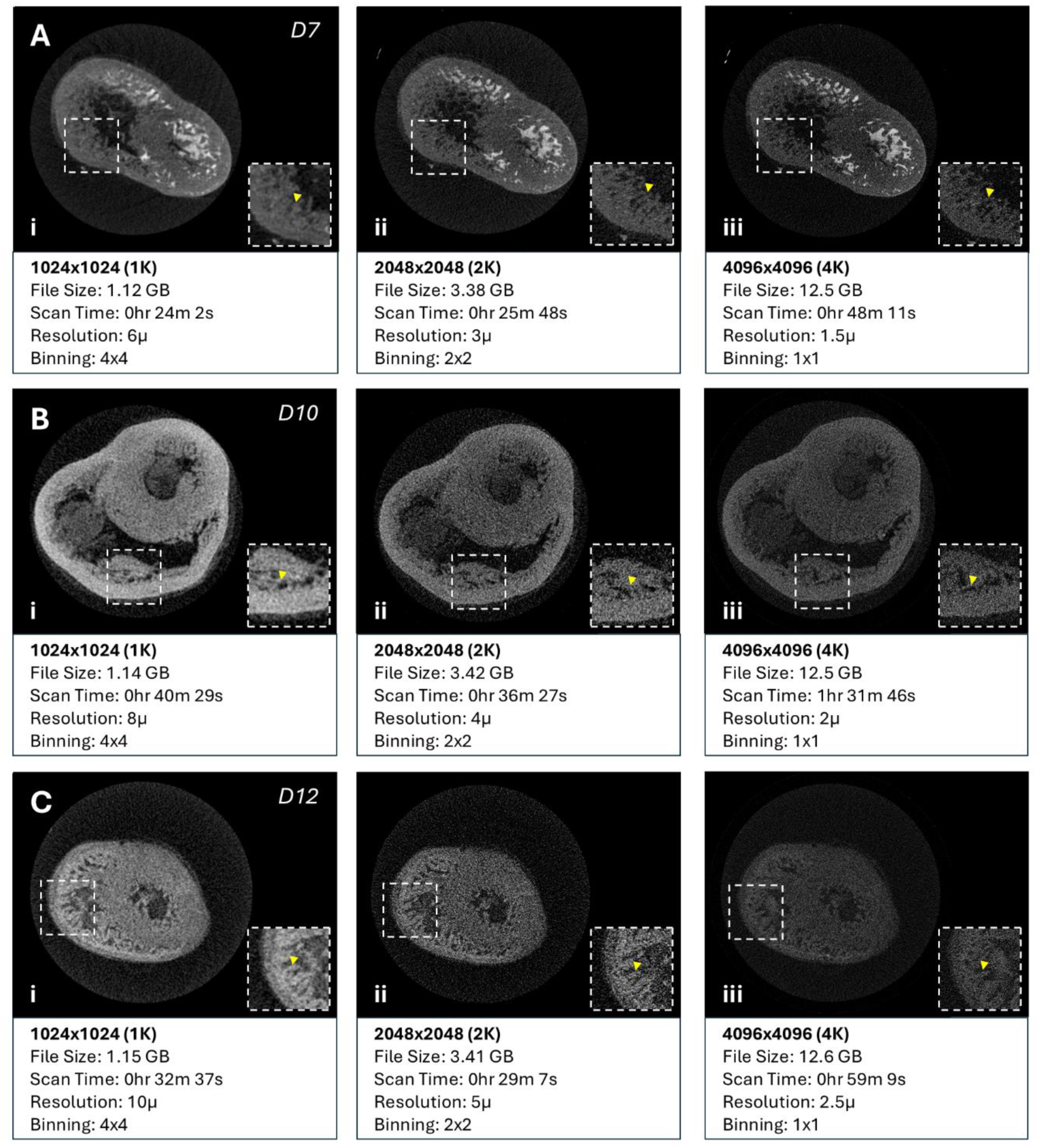
Selection of camera binning. **A-C:** Reconstructed transverse cross-sections from (**A**) day 7 (D7), (**B**) day 10 (D10), and (**C**) day 12 (D12) embryonic hearts scanned using a (**i**) 1K camera (4x4 binning), (**ii**) 2K camera (2x2 binning), and (**iii**) 4K camera (1x1 binning). Yellow arrows point to trabeculae.

On the Bruker SkyScan 1272 micro-CT unit, pixel binning is not performed in the traditional sense, instead the X-ray detector has 4 camera modes, (512x512px, 1024x1024px, 2048x2048px, and 4096x4096px), these are cropped detector modes. As the 4K (4096x4096px) detector uses the full range of the detector, more pixels are acquired, producing a smaller effective pixel size, consequently, it is more sensitive to ring-artefacts, motion misalignment and noise (Fig 4. iii) (Kyriakou et al., 2009), whilst also increasing the time taken per projection which substantially increases the total scan time (Ketcham & Carlson, 2001).

To establish the best camera binning, samples at each timepoint were scanned using the 1K, 2K or 4K camera to establish whether increasing the camera binning had any significant dampening of image quality. Increasing binning from 1x1 (Fig 4. iii) to 2x2 (Fig 4. ii) reduced scan time whilst maintaining sufficient image quality to resolve anatomical structures of interest. Further increasing binning to 4x4 (Fig 4. i) resulted in a noticeable loss on image detail which resulted in difficulty delineating tissue boundaries and trabeculae (as represented by the yellow arrows in Figure 4). There was also no noticeable difference in scan duration between the 2x2 (Fig 4. Bii-Cii) and 1x1 binning (Fig 4. Biii-Ciii), based on these comparisons the 2x2 binning parameter (2K) was selected. The 2x2 binning scan time for day 10 was 36 minutes (Fig 4. Bii) compared to 1 hour 31 minutes during the 1x1 binning scan (Fig 4. Biii), as there was no significant increase in scan quality, the 2x2 binning (2048x2048px) was selected for the best balance between image quality and acquisition time. Higher isotropic resolution does not always improve biological accuracy, often signal-to-noise ratios and artefact levels need to be considered (Mizutani & Suzuki, 2012). As a compromise, the 2K camera binning option (Fig 4. ii) was chosen to maintain high resolution, lower scanning time and fewer artefacts.

### Optimal scanning parameters identified for each timepoint

The optimal scanning parameters (isotropic voxel size, filter, camera binning, rotational step and frame averaging) across each of the developmental timepoints were obtained. Table 2 lists optimised image resolution and total scanning time, whilst still obtaining high-resolution 3D renders for downstream quantitative and morphometric analysis.

**Table 2.** Optimal scanning parameters for each individual timepoint.

| SkyScan 1272 | D7 | D10 | D12 |
| --- | --- | --- | --- |
| Isotropic Voxel Size | 3-4 μm | 4-5 μm | 5-6 μm |
| X-ray Power | 35kV-200μA | 45kV-200μA | 45kV-200μA |
| Filter Selection | No Filter | Al 0.25 | Al 0.25 |
| Camera Binning | 2048x2048 | 2048x2048 | 2048x2048 |
| Rotational Step | 0.5° | 0.5° | 0.5° |
| Averaging Frames | 3 | 3 | 3 |
| Scan Time | 29m 15s | 31m 29s | 31m 29s |
| Rotation | 180° | 180° | 180° |

The only change in parameter between time points is the filter selection used. The specimen size of the D10 and D12 hearts required use of the 0.25mm aluminium filter to correct x-ray attenuation as shown previously in Figure 3. Voltage and current were automatically adjusted by the Bruker SkyScan 1272 when changing between filters and voxel size.

### Determination of reconstruction parameters and pre-analysis operations

When reconstructing micro-CT datasets, it should be considered which parameters are used to obtain a high-quality 3D render with minimal artefacts (Vickerton et al., 2013). As the cardiac soft tissue contrast was artificially enhanced using buffered Lugol’s iodine, staining intensity varied across specimens, likely due to uneven penetration into each sample potentially due to biological heterogeneity and diffusion kinetics. Iodine leaching into the PBS used to load the sample into the PCR tubes prior to scanning can also affect attenuation of x-rays penetrating through the sample (Bouxsein et al., 2010; Pauwels et al., 2013).

It is also imperative to consider where to set the global greyscale threshold to allow standardised comparisons across biological specimens within a dataset, this should be done post-scanning optimisation. Our approach included decreasing the higher density threshold range to remove the signal from the PCR tube from all the samples and increasing the lower density threshold to remove air and PBS from the background of the sample to maximise the greyscale range allocated to the biological structures present (Rovaris et al., 2018).

### Ring-artefact corrections

Ring-artefacts are a common post-reconstruction artefact that can be removed with the ring-artefact correction (RAC) tool during the reconstruction process in NRecon. Ring-artefacts are circular streaks observed across the reconstructed dataset. Ring-artefacts need to be corrected as they interfere with global thresholding by artificially altering the greyscale intensities within each sample (Sijbers & Postnov, 2004).

Artefact correction levels of RAC were demonstrated in Figure 5 using NRecon. RAC correction is conducive to remove artefacts as they can impair quantitative 3D analysis, but an overcorrection can cause blurring and additional artefacts (Gaêta-Araujo et al., 2020). Studies have shown that ring artefacts can pose significant challenges for quantitative analysis, therefore it is important to ensure the artefacts have been corrected prior to volumetric analysis (Sijbers & Postnov, 2004; Hasan et al., 2010). Beam-hardening artefacts are also common in mineralised tissue, less common in soft tissue. Beam-hardening artefacts were minimal within the biological specimens due to relatively uniform attenuation across the specimen, however, some specimens contained a whitening of the PCR tube, where a correction of 28% was applied in NRecon (Meganck et al., 2009; Vásárhelyi et al., 2020).

**Figure 5.**
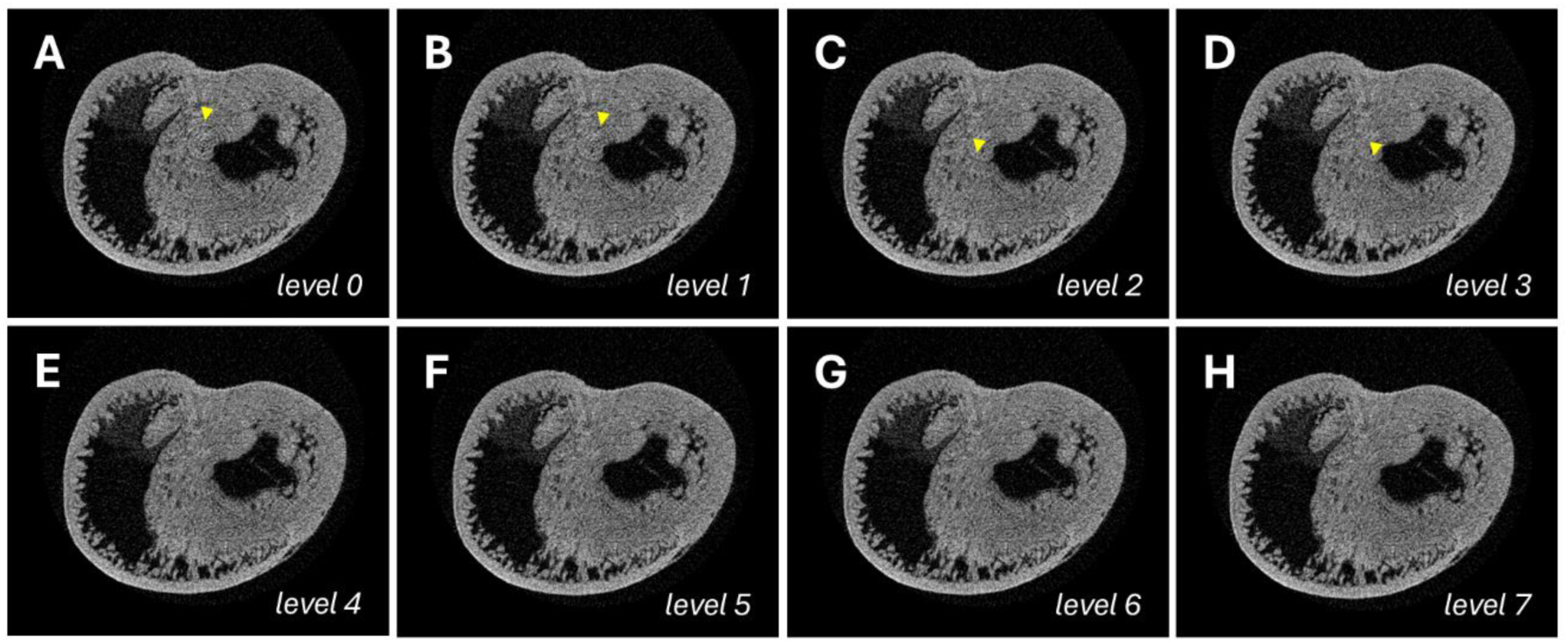
Ring artefact correction (RAC) tool. Transverse cross-section of an embryonic day 12 chicken heart in NRecon with (**A**) no ring-artefact correction applied increasing by 1 to (**H**) with a ring-artefact correction of 7. Yellow arrow indicates ring artefacts.

Ring-artefacts were present with no correction applied. With a correction power of 4 (Fig 5. E), the artefact was almost completely removed. A correction of 5 (Fig 5. F) had no visual improvement over a RAC of 4. It was determined that a RAC of 4 did not affect tissue structure or quality of individual reconstructions within the dataset.

### Re-orientation of biological specimens

Datasets were standardised and orientated to the correct alignment for consistent analysis across specimens. DataViewer was used to re-align each specimen so that the frontal face of the heart was positioned anteriorly in the coronal plane. A standardised orientation preserves the anatomical axis and prevents orientation-related variability in post-reconstruction volumetric analysis.

Figure 6 is an of an embryonic day 10 specimen being re-orientated post micro-CT acquisition. This approach was replicated in samples collected from embryonic days 7 and 12.

**Figure 6.**
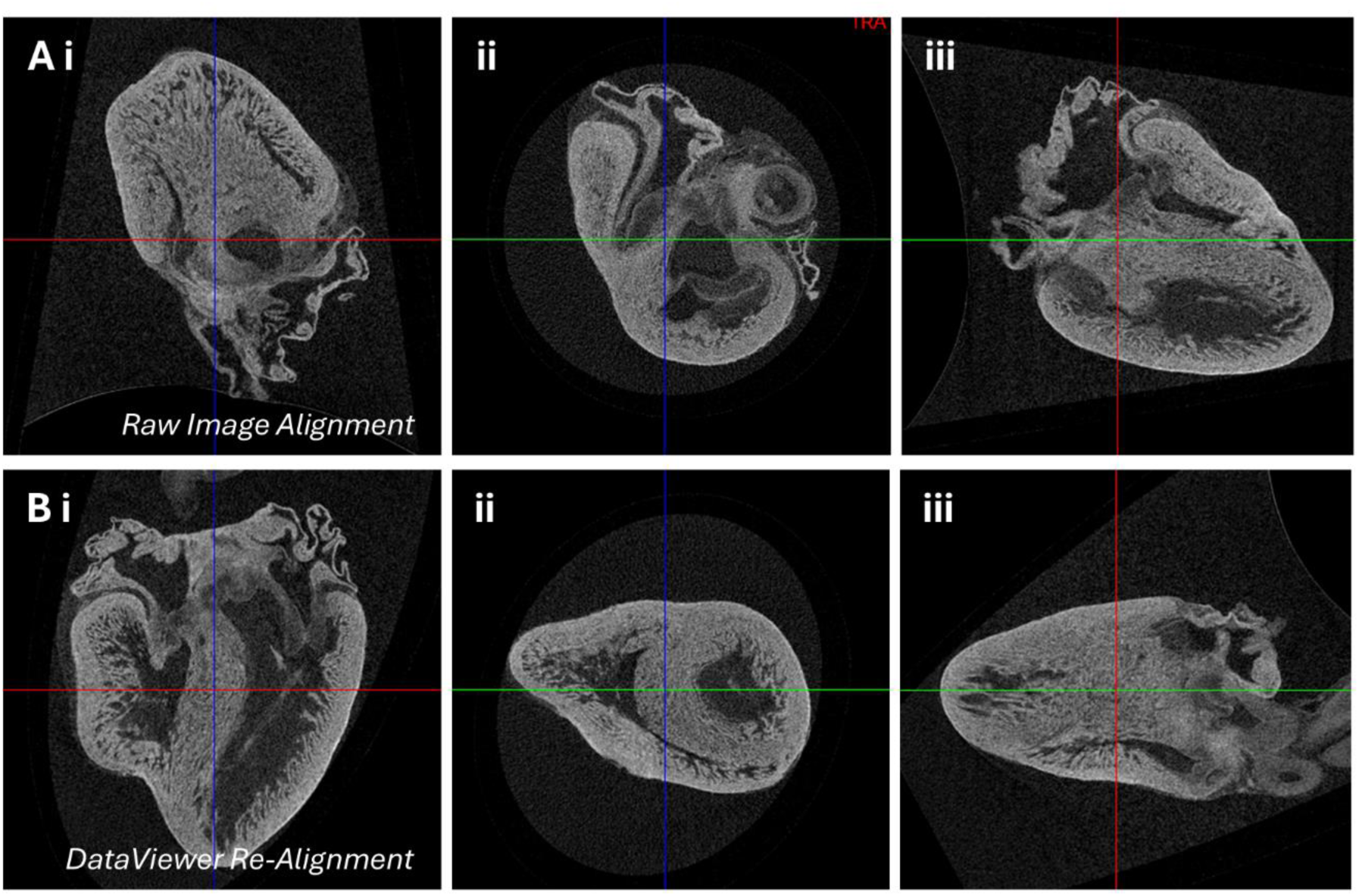
DataViewer re-alignment tool. **A:** Unprocessed image of a D10 chicken heart post reconstruction. **B:** Post-realignment using DataViewer to manipulate sample orientation for standardised analysis. Images viewed in (**i**) coronal, (**ii**) transverse and (**iii**) sagittal optical planes. The red line indicates the transverse plane, the blue line indicates the sagittal plane, and the green line indicates the coronal plane.

### Optimal reconstruction parameters in NRecon

The optimal reconstruction parameters (global thresholding and smoothing) across each of the developmental timepoints were obtained (Table 3). RAC and beam hardening correction tools were performed on a sample-by-sample basis as each sample requires a different level of correction.

**Table 3.** Reconstruction parameters for each individual timepoint.

| NRecon | D7 | D10 | D12 |
| --- | --- | --- | --- |
| Global Thresholding | 0.04-0.2500 | 0.01-0.1000 | 0.02-0.0850 |
| Smoothing | Gaussian, 1 | Gaussian, 1 | Gaussian, 1 |

### Post-processing operations

The use of smoothing operations such as gaussian or median filtering are commonly used and widely accepted to reduce noise for quantitative assessment (Shahmoradi et al., 2016). Gaussian smoothing applies a weighted distance-dependent blur where neighbouring pixels contribute more than distant pixels. This method is effective at reducing high-frequency noise whilst preserving anatomical gradients (Shahmoradi et al., 2016). Median filtering replaces individual pixels with the median of its neighbouring pixels; this method is particularly effective at removing salt-and-pepper noise (Shahmoradi et al., 2016). For this study, a mild gaussian smoothing value of 1 was chosen to conservatively reduce high-frequency noise from the detector to preserve anatomical boundaries prior to further processing.

Smoothing was not optimised during the reconstruction process in NRecon, instead focusing on the post-processing tools within Dragonfly. A post-processing tool operation task list consisted of a median filter (3D; kernel - 3) to smooth salt-and-pepper noise (Gau & Liu, 2015). This was followed by a morphological closing operation (2D; kernel - 3) to reduce background noise whilst preserving tissue morphology as shown in Figure 7B (de Albuquerque et al., 2020).

**Figure 7.**
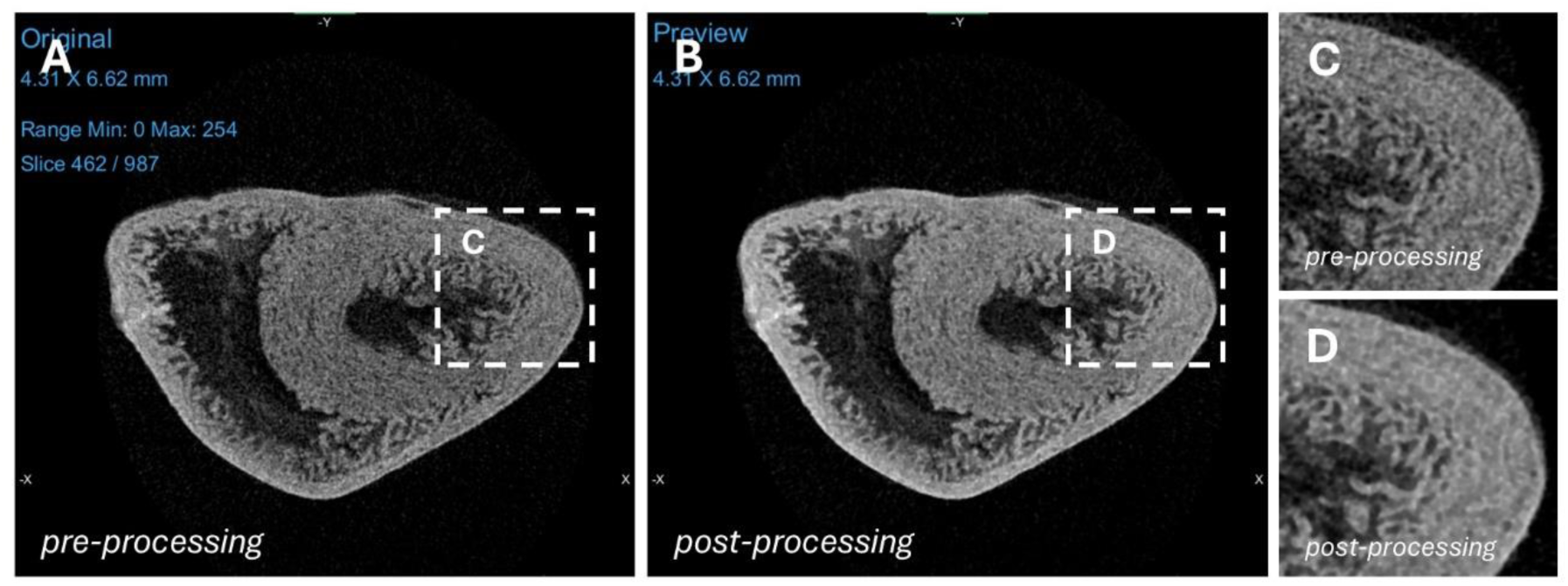
Post-reconstruction processing operations in Dragonfly. **A-B:** Post-reconstruction operations In Dragonfly showing the radiograph (**A**) before processing and (**B**) after processing of a transverse heart section. **C-D:** Enlarged radiographs of (**C**) pre-processing and (**D**) post-processing.

The smoothing and closing operations were highly effective in reducing image noise. Prior to processing (Fig 7. C), the micro-CT images appeared grainy, making the trabecular boundaries less distinct and more difficult to segment accurately. Following these post-processing operations (Fig 7. D), image noise was reduced, the trabeculae became more clearly defined, and segmentation was more accurate and consistent.

Post-processing thresholding is a method of segmentation to separating the greyscale range of pixels into the foreground and background (Buyuksunger et al., 2024). Removing noise using a median smoothing filter is a standard approach to remove stochastic detector noise whilst preserving tissue and anatomical boundaries for consistent and reproducible segmentation across biological specimens (Aslam et al., 2023; Gau & Liu, 2015)

The task list for segmentation in Dragonfly included pre-processing where a median smoothing filter (3D; 3x3x3 kernel) was applied to reduce noise, followed by a morphological closing (2D; 3x3 kernel) to remove small-speckle artefacts during thresholding. Intensity-based threshold selection used the brush tool to select the regions of interest. To calculate wall volume, 28-connected components were removed to eliminate small non-anatomical structures that were incorrectly selected during the segmentation. For lumen volume, the 6-connected components option was selected to retain only the primary lumen cavity, and all non-connected structures were removed from the segmentation.

### Volumetric analysis using Dragonfly

As the aim of this study was to calculate both ventricular wall and lumen volume. Dragonfly was selected due to its segmentation tools to permit user-friendly delineation of regions of interest (Buyuksunger et al., 2024; Dragonfly, n.d.-1; Dragonfly, n.d.-b).

As there was normal and expected biological variation in cardiac morphology between samples, data was normalised prior to interpretation and comparison. Wall measurements (Fig 8. A) and lumen measurements (Fig 8. B) were taken from 6 hearts harvested from day 10 embryos.

**Figure 8.**
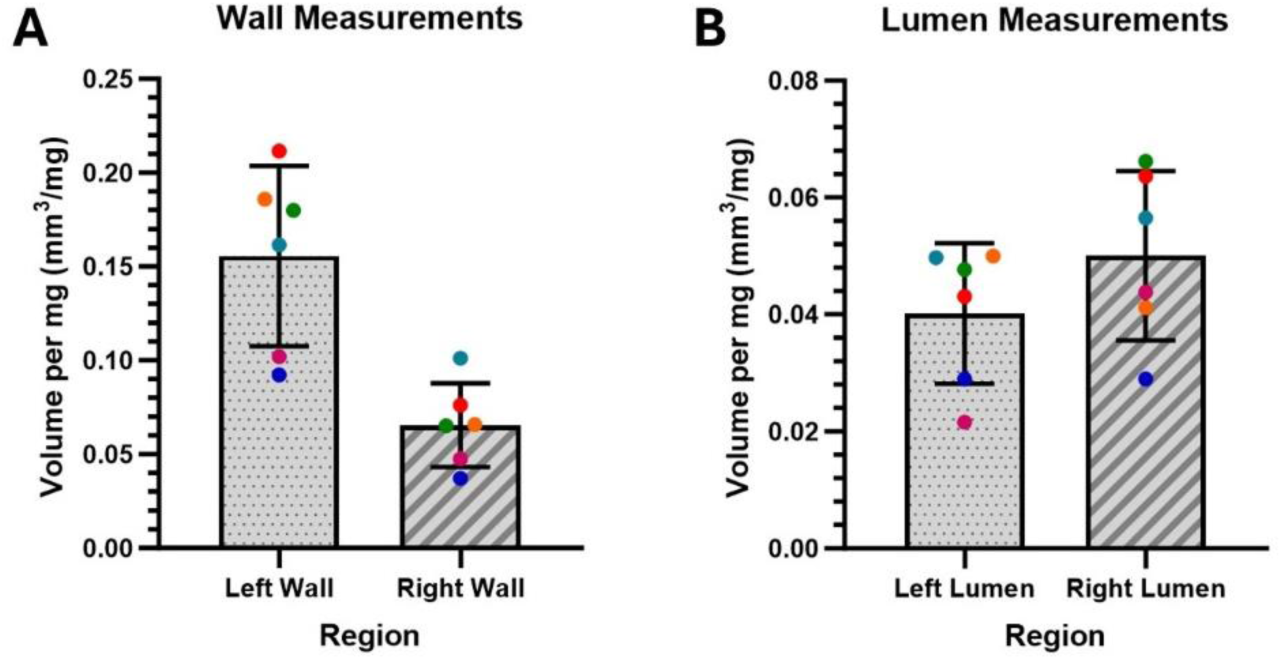
Volumetric myocardial and luminal data collected from Dragonfly. **A:** Quantification of myocardial wall volume in the left (0.16±0.05 mm^3^) and right (0.07±0.02 mm^3^) ventricles of a day 10 heart, plotted as individual biological replicates (n=6). **B:** Quantification of left (0.04±0.01 mm^3^) and right (0.05±0.01 mm^3^) luminal volume of a day 10 heart, shown with individual biological replicates (n=6). Each colour represents a separate biological replicate.

The volumetric measurements collected from the ventricular wall (Fig 8. A) and lumen (Fig 8. B) were consistent indicating low inter-embryo variability at the embryonic day 10 timepoint, suggesting that micro-CT acquisition and segmentation produced reproducible morphometric data outputs. The small standard deviation within each category from the LV wall (0.16±0.05 mm^3^) and RV wall (0.07±0.02 mm^3^) demonstrates that the segmentation is consistent and can reliably extract volumetric features across biological replicates. These measurements are consistent with previous descriptions indicating that the left ventricular myocardial volume is greater than right ventricular myocardial volume (Faber et al., 2021).This concordance demonstrates the ability of micro-CT to deliver accurate, high fidelity volumetric morphometry of embryonic cardiac structures.

### Reversible contrast-stain and downstream imaging

#### Reversible-Lugol stain

We assessed the compatibility of reversible Lugol’s contrast staining with downstream histological and immunohistochemical workflows. As a well-established micro-CT contrast agent, Lugol’s enables non-destructive three-dimensional visualisation of specimens before conventional destructive analyses, including histopathology (Hopkins et al., 2015; Metscher, 2009). Iodine was chosen over other contrast agents such as phosphotungstic acid as it can be readily de-stained from samples using sodium thiosulphate (Hopkins et al., 2015; Metscher, 2009), increasing the likelihood of antigen epitopes are retained for downstream immunohistochemistry. Although Hopkins et al. (2015) demonstrated that iodine can be removed with sodium thiosulphate to allow subsequent histology and immunostaining in adult peripheral nerve tissue, their evaluation was purely qualitative. This study expanded on this by applying reversible Lugol’s staining to enable quantitative volumetric analysis of embryonic cardiac tissue, including chamber-specific wall and lumen measurements that are directly relevant to developmental morphogenesis and CHD aetiology (Fig 8).

Following micro-CT imaging, Lugol’s stain was removed by incubating the samples in 2.5% sodium thiosulfate for 24 hours (Fig 9. iii) before FFPE processing and downstream immunostaining. Sodium Thiosulphate is a well-established reducing agent for the removal of iodine-based stains (Asakai & Hioki, 2011; Lanzetti & Ekdale, 2021), although low concentrations (below 10% w/v) are recommended to minimise alterations to fixed tissue specimens (Gignac et al., 2016). Destaining was effective, with destained tissue (Fig 9. iii) closely resembling unstained tissue morphology (Fig 9. i). Furthermore, no discernible shrinkage after destaining was observed in day 10 hearts, with sample diameter only changing from approximately 1.45mm before staining to 1.42mm after destaining.

**Figure 9.**
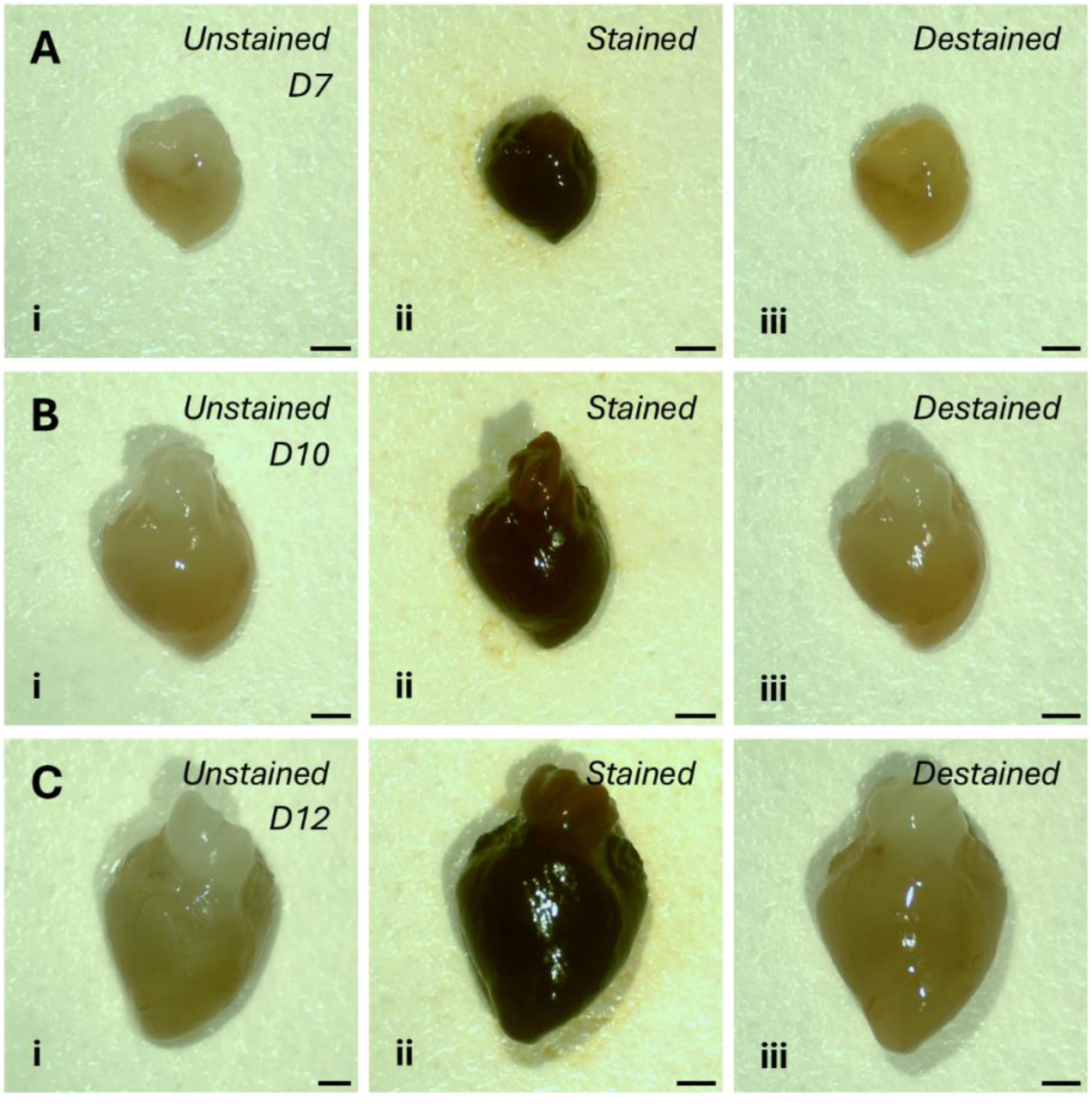
Images of embryonic chicken hearts pre- and post-staining and post-destaining. **A-C:** Embryonic hearts at (**A**) day 7, (**B**) day 10, and (**C**) day 12. Specimens were imaged (**i**) prior to staining, (**ii**) after staining, and (**iii**) after de-staining. Scale bar is 500μm.

#### Multimodal workflow and subsequent histopathology

To assess the compatibility of iodine-enhanced micro-CT specimens with downstream histological analysis, samples were probed for phosphohistone-H3 (PHH3) as a nuclear-specific antibody and β-actin as a cytoplasmic-specific antibody alongside wheat germ-agglutinin (WGA) as a non-antibody based staining technique which targets glycoproteins on the cell membrane to determine whether iodine had interfered with antigenicity and lipid stability as shown in Figure 10A-B (Ryva et al., 2019). H&E histological staining was also performed (Fig 10. C-F) to identify compatibility with broadly used traditional techniques.

**Figure 10.**
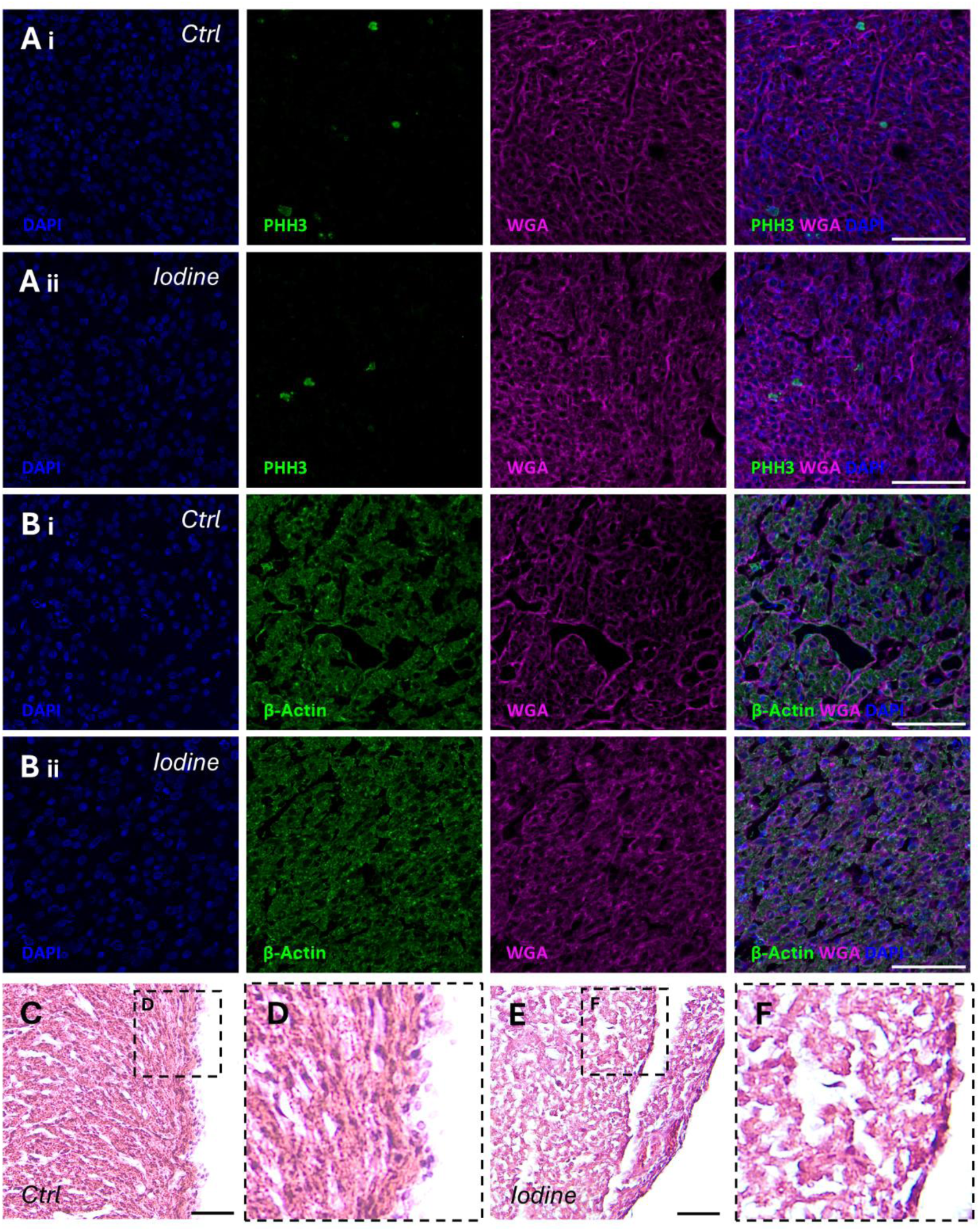
Immunofluorescent and histological staining of cardiac tissue. **A-B:** Immunofluorescent micrographs (40X objective) taken using a Zeiss LSM800 confocal microscope of (**A**) PHH3 (green), WGA (magenta), and DAPI (blue); and (**B**) β-actin (green), WGA (magenta) and DAPI (blue) in (**i**) control and (**ii**) iodine-destained samples. **C-F:** Histological H&E-stained tissue sections from (**C-D**) control and (**E-F**) iodine-destained samples imaged using a brightfield microscope (20X objective). Scale bar is 50um.

Initial attempts at paraffin-sectioning at 5μm resulted in tissue fragility and crumbling during microtomy, suggesting that the tissue became more brittle after iodine-staining. Sections were subsequently prepared at 10μm thickness which improved tissue integrity and allowed reliable downstream staining and imaging. Destained samples presented with similar fluorescent intensity and staining profile (Fig 10. Aii-Bii) compared with non-iodine-stained controls (Fig 10. Ai-Bi). In addition to the antibody-based labelling, WGA staining visualised distinct cellular membranes, suggesting that iodine destaining did not affect tissue architecture and demonstrating that lectin-based recognition sites are retained.

#### Assessment by Brightfield and Fluorescent Microscopy Techniques

Traditional histological staining techniques (H&E) were used to visualise cellular architecture. We found staining intensity of the nuclei appeared reduced in the iodine-destained sample’s (n=3) (Fig 10. F) compared with the control (Fig 10. D). This resulted in less contrast between the nuclei (haematoxylin stain) and cytoplasm (eosin stain) and was observed across multiple biological specimens and technical repeats (n=3). Although nuclei staining provided limited nuclear contrast following the de-staining process, cytoplasmic staining remained well preserved, allowing overall assessment of tissue morphology and architecture. However, reduced nuclear staining may limit analysis that rely on accurate delineation and visualisation of cell nuclei, such as nuclear morphology and cell density assessment. Importantly, successful PHH3 immunostaining demonstrates that reduced nuclear staining using traditional haematoxylin does not necessarily reflect a loss of antigen availability, indicating that nuclear protein localisation and proliferative/cell density assessment remain feasible using other techniques.

Positive immunostaining for PHH3 and β-actin (Fig 10. Aii-Bii) indicates that both nuclear and cytoplasmic antigens remain detectable following iodine-enhancement and subsequent sodium thiosulphate destaining, enabling the assessment of proliferative index and cytoskeleton architecture. Positive WGA staining also demonstrates that the multimodal workflow is not limited to antibody-based staining techniques and that lectin-based techniques are also compatible, enabling visualisation of cell membranes and assessment of cross-sectional cell area. However, very little information is available of which antigens are preserved following iodine staining and subsequent destaining procedures, we recommend user-specific optimisation of desired antibodies.

Overall, our optimised protocol holds several distinct advantages; resolution and contrast restraints inherent to micro-CT mean that it cannot provide cellular-level information (Jiang et al., 2005). By integrating alongside the traditional 2D workflow, it was possible to non-destructively visualise gross structure, allowing screening for gross morphological abnormalities in cardiac structure (e.g. congenital defects and other regions of interest) prior to the destructive sectioning and imaging of valuable specimens for cell and molecular spatial studies. These complementary modalities are particularly advantageous in the field of developmental biology; the visualisation of transient and continually remodelled structures alongside concurrent molecular information in the same samples can provide insights into disease pathophysiology which may be masked by intra-embryo variability. For example, ventricular septal defects (VSD) arise during a narrow developmental window, and embryos of the same stage or treatment may differ in whether a defect is present. This workflow allows a VSD to be identified confidently in the micro-CT dataset and then enables probing of molecular-level changes within that exact embryo, which is not possible with traditional destructive workflows that require separate specimens for morphology and molecular analysis (Williams et al., 2019).

#### Limitations

This study demonstrates that fixed *ex-vivo* embryonic hearts can be used to quantify whole-organ geometry, luminal volume and phenotypic characteristics through a multimodal approach integrating micro-CT with downstream immunohistochemistry. As with many *ex-vivo* approaches using fixed tissue, this methodology does not capture the dynamic physiological parameters such as intracardiac pressure, contractility, or flow rates. Consequently, the volumetric measurements reflect structural morphology rather than functional haemodynamic performance.

### Conclusions

We have shown that micro-CT is a useful high-resolution imaging tool to visualise embryonic chick cardiac tissue and that volumetric analysis and segmentation of the cardiac macro-structure is robust and reproducible, allowing analysis of individual regions of interest within the heart, whilst also preserving the fine structure of the valves and trabeculae. Optimised scanning and reconstruction parameters enabled high-resolution 3-D modelling of embryonic hearts at multiple developmental timepoints, demonstrating that this approach may be valuable for investigating cardiac hypertrophies, myopathies and stenosis throughout embryonic development. Importantly, the reversible Lugol’s based enhancement stain preserved antigen availability to enable compartmental staining of cytoplasmic and nuclear protein targets within the same specimen.

This dual-modality approach combines whole-organ 3D morphology study with simultaneous targeted cellular and molecular analysis in the same sample, a key advantage when researching CHDs due to incomplete penetrance of suspected defects. Although this study specifically focused on embryonic chicken tissue in the cardiac context, we strongly suspect that this workflow will prove broadly applicable to other small tissues and organs where combining structural context with molecular assays is beneficial, including alternative small animal models (mouse, rat and zebrafish), as well as non-cardiac contexts such as characterisation of tumour structure and bioengineered tissue constructs.

## ACKNOWLEDGEMENTS

We would like to thank the Sheffield Multimodal Imaging Centre for access to the micro-CT imaging facility, which was initially funded by the European Regional Development Fund, grant number 28R19P03971). We would also like to thank Dr Lucy Dascombe and Olivia Doolan for providing extensive micro-CT training and technical support. Data collection was performed using Dragonfly 3D World, provided by Comet Technologies Canada Inc., Montreal, Canada. We would also like to thank Prof Marysia Placzek at the Bateson Centre (University of Sheffield) for providing the eggs used during this study.

## CONTRIBUTIONS

JD, PS, LR, and MH contributed to the conception, design, acquisition and interpretation of the data and the drafting/revising of the manuscript. NA contributed to the experimental design, interpretation of data, training on micro-CT and revising of the manuscript.

